# Estimating anisotropy of single cells using a compliant biaxial stretcher

**DOI:** 10.1101/2025.06.27.661720

**Authors:** Himanshu Marwah, Hetarth Bhatt, Neeraj Fartyal, Rohit Nautiyal, Sukh Veer, Pramod A Pullarkat, Sreenath Balakrishnan

**Affiliations:** School of Mechanical Sciences, Indian Institute of Technology Goa; School of Interdisciplinary Life Sciences, Indian Institute of Technology Goa; Raman Research Institute, Bengaluru, India

**Keywords:** biaxial stretching, compliant mechanism, cell manipulation, mechanical characterization, micro-robotics

## Abstract

Understanding the mechanical properties of single cells is essential for elucidating their responses to mechanical cues and plays a critical role in development, disease progression, and mechanotransduction. However, probing these properties at the single-cell level remains technically challenging. Traditional methods for probing cellular mechanics are often restricted to a single deformation mode and rely on complex instrumentation, limiting their adaptability. Hence, no single platform currently offers multi-modal cell manipulation, which motivates the development of more versatile tools. Towards this, we are developing an ensemble of compliant micromechanisms that enables diverse modes of cell manipulation on a single chip. Building on our earlier work with a uniaxial stretcher, here we demonstrate estimation of cellular anisotropy using a biaxial stretcher. Since the mechanism transforms motion using deformation, cell stretching forces can be estimated through image-based displacement measurements, without additional sensing modalities, considerably simplifying the device fabrication and operation. By assembling these uniaxial and biaxial stretching micromechanisms on a single chip, we can establish a multi-modal manipulation platform for single cells, which can be augmented with shearing and twisting modes in the future. Such a simple and versatile platform presents a paradigm shift in mechanical testing of single cells.

## 1 Introduction

Mechanical cues in the cellular microenvironment play a central role in regulating various biological processes, including embryogenesis, stem cell fate decisions, and cancer progression [1, 2]. These cues influence cellular behavior and provide insight into the mechanical state and health of tissues. Understanding how cells sense and respond to such mechanical stimuli—broadly referred to as mechanotransduction—requires tools capable of applying and measuring forces at the single-cell level. Towards this, several experimental techniques have been developed to probe the mechanical properties of single cells. Atomic force microscopy (AFM) characterizes stiffness by indenting the cell surface using a cantilever-mounted tip [1]. Micropipette aspiration applies suction to deform the cell membrane [2], while optical tweezers (OT) use focused laser beams to trap and manipulate beads attached to the cell surface [3]. In parallel-plate rheometer (PPR), a cell is attached between two plates and stretched (or compressed) by moving one of the plates [4]. Other approaches include magnetic twisting cytometry (MTC), which induces shear via magnetically actuated beads [5], and substrate stretchers, which stretch the substrate beneath the cell [6]. Despite their effectiveness, each of these techniques are typically limited to a single mode of deformation and require distinct, often unwieldy, setups for each manipulation mode. In an intriguing study published recently, several single-cell manipulation techniques such as AFM, MTC, PPR, and OT were used to measure the mechanical properties of MCF7 cells [7]. Due to the varied methodology and cell states during measurements, such as adhered to a substrate in case of AFM, MTC and PPR and suspended in the media case of OT, the elastic modulus estimated by each of these techniques showed large variability, to the tune of a few orders of magnitude. Further, the specialized nature of these techniques necessitated contributions from several research groups with each group typically contributing only one measurement technique. This study highlights the need for a unified, convenient framework for single-cell manipulation that can perform a diverse set of manipulation modes. We posit that such a framework can be realized using monolithic micromechanisms, without joints, designed using the compliant mechanism design methodology.

Compliant mechanisms generate motion through elastic deformation rather than hinges or sliding joints, making them suitable for integration into a lab-on-a-chip. Their monolithic design allows multiple mechanisms—each tailored to a specific mode of deformation-to be fabricated and operated independently. Schematics for four compliant mechanisms for uniaxial and biaxial stretching, twisting and shearing single cells are shown in Fig 1. Uniaxial stretching is achieved using a folded-beam structure that pulls the cell along one axis (Fig. 1A). In Fig. 1B, the cell is attached to four triangular platforms arranged in orthogonal directions. When actuated at a single input point, the mechanism generates symmetric displacements in both horizontal and vertical directions, resulting in biaxial stretching. In Fig. 1C, the cell is attached between an overhanging gripper at the top and the substrate at the bottom. By rotating the gripper, the cell is twisted along the axis perpendicular to the plane. In Fig. 1D, the cell is anchored between two parallel beams. One of these beams is laterally displaced while the other remains fixed, inducing shear. All these mechanisms can be microfabricated and mounted on a coverslip, thereby enabling high-resolution imaging during manipulation. Among these schematics, we have previously demonstrated uniaxial [8, 9] and biaxial stretching [10, 11, 12] whereas the twisting and shearing mechanisms are concepts presented to highlight the versatility of this framework.

**Figure 1:**
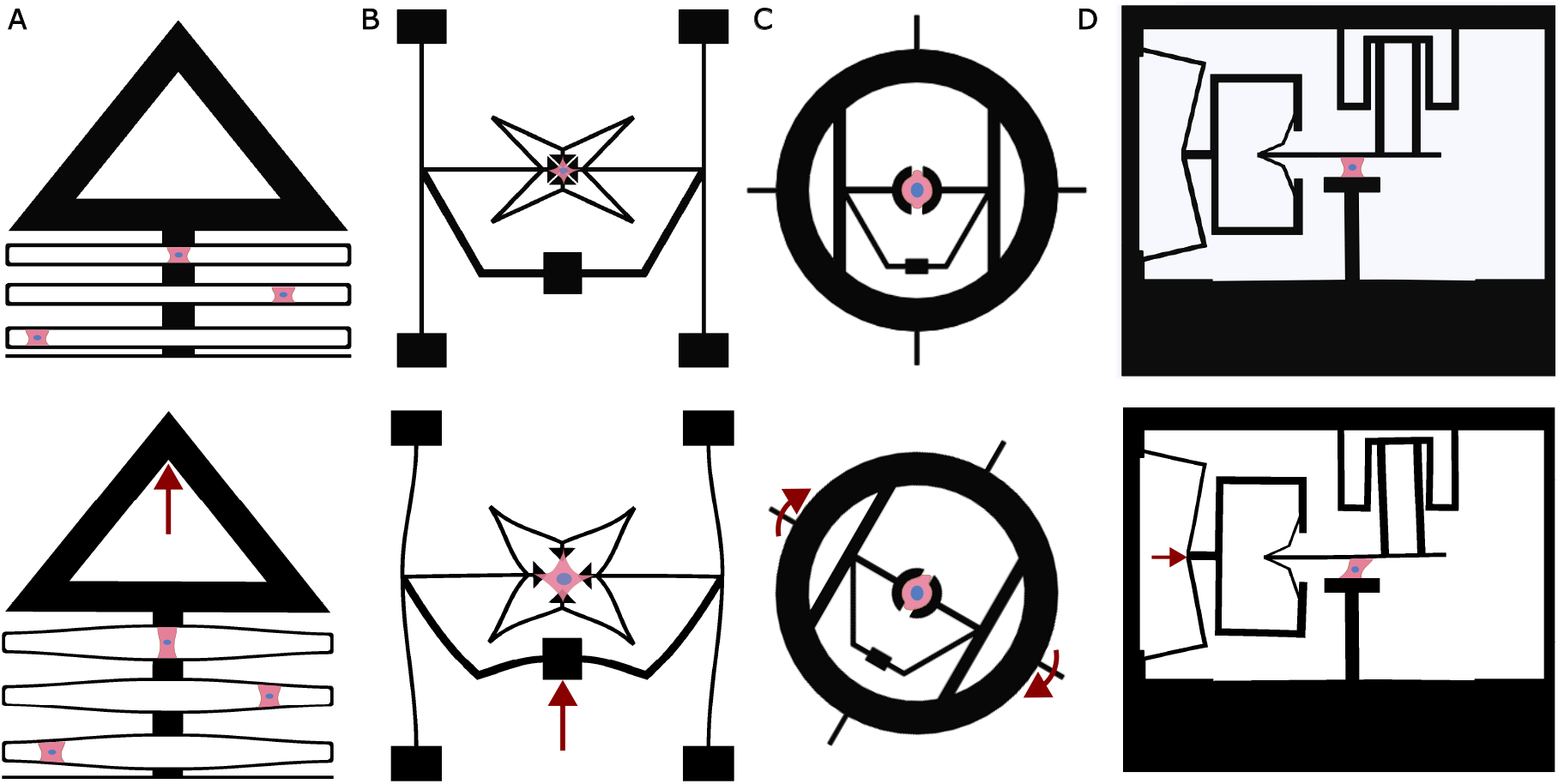
Single-cell mechanical manipulation using compliant mechanisms. Top row: undeformed (initial) configurations of the compliant mechanisms with adhered cells. Bottom row: deformed configurations under actuation. Red arrows indicate the actuation applied to the compliant mechanisms. (A) Uniaxial stretching, (B) Biaxial stretching, (C) Twisting, and (D) Shearing

In the uniaxial stretcher, the force applied to the cell could be estimated from the mechanism’s deformation [9]. However for the biaxial stretcher, while single cells could be stretched biaxially[12], the stretching force could not be estimated, and therefore cellular anisotropy could not be characterized. In the present study, we address this limitation by enabling quantitative mechanical characterization using the biaxial stretcher. We derive closed-form expressions that relate mechanism displacements to cell stretching forces using Euler–Bernoulli beam theory and Castigliano’s theorems. We validate these expressions both at the macro and micro scales. At the macro scale, we fabricated prototypes made out of acrylic to stretch calibrated extension springs and polydimethylsiloxane (PDMS) membranes with known mechanical properties. Micro scale devices are fabricated using SU-8 photoresist using a two-layer lithography process. The Young’s modulus of SU-8 was estimated by actuating this micromechanism using an optical fiber-based force transducer. Finally, using the micromechanism we stretch individual U87-MG glioblastoma cells biaxially. Cell strains are measured from the microscopy images and cell stresses are computed from the mechanism deformation. These data are fit to a 2D Fung model to extract mechanical parameters. The results reveal direction-dependent stiffness and coupling between orthogonal directions, indicating cellular anisotropy. This work demonstrates a sensor-free, image-based method for quantifying single-cell anisotropy and lays the groundwork for an integrated, multi-modal single-cell manipulation platform.

## 2 MATERIALS AND METHODS

### 2.1 Design and microfabrication of biaxial stretching mechanism

Our initial design for a biaxial stretching device was based on a geometric configuration using a re-entrant shape (Fig. 2A), as described in earlier work [10]. Re-entrant geometries contain internal angles greater than 180°, forming concave polygons. These shapes, and their three-dimensional equivalents, re-entrant polyhedra, exhibit auxetic behavior or negative Poisson’s ratio [13]. When stretched along one axis, they expand in the perpendicular direction, enabling biaxial extension.

**Figure 2:**
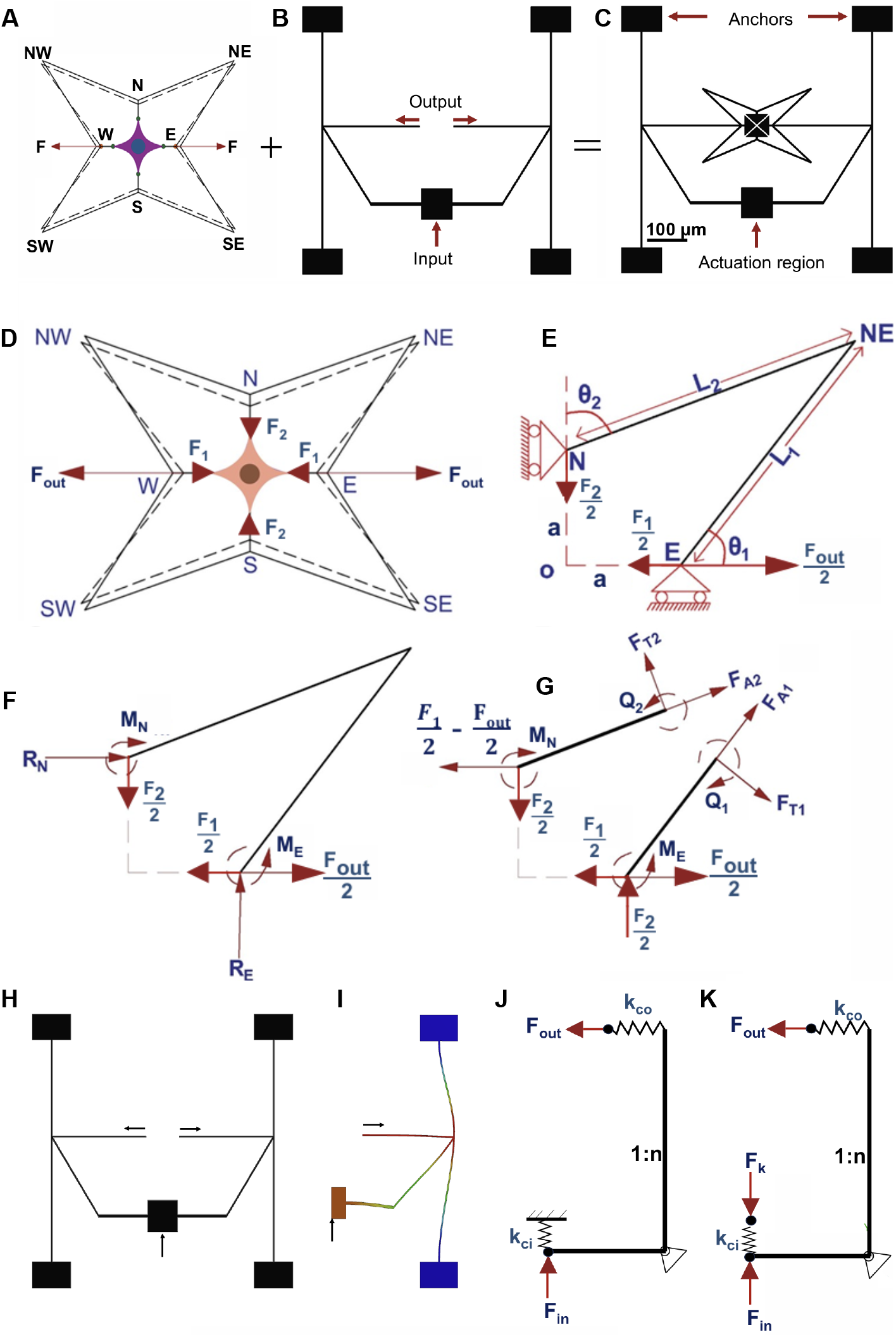
Mechanism design. (A) Double-input biaxial stretcher based on re-entrant structures. A single cell is attached between N (north), S (south), E (east), and W (west) points. (B) Single-input, double-output gripper. (C) Single-input biaxial stretcher obtained by combining the re-entrant shape (double-input) with the gripper (single-input, double-output). (D) Two-input biaxial stretching auxetic mechanism. (E) The analysis can be reduced to one-quarter of the design with symmetric boundary conditions. (F) Forces applied on the quarter design. (G) Free body diagram of the quarter design. Spring-leverage model (SL): (H) a compliant gripper, (I) its symmetric half used in deformation analysis, (J) its representation as a lever with geometric amplification factor *n*, input-side spring stiffness *k*_*ci*_, and output-side spring stiffness *k*_*co*_, and (K) Free body diagram of the SL model.

To actuate the mechanism, equal and opposite forces (F) were applied along the horizontal axis at the east (E) and west (W) points. The geometric of the re-entrant structure is designed such that the outward displacement at the north (N) and south (S) and east (E) and west (W) points are equal thereby achieving equibiaxial stretching [10]. To eliminate the need for dual forces, a compliant gripper mechanism was integrated to convert a single input into two opposing horizontal outputs (Fig. 2B). These outputs were directed to the E and W points of the re-entrant structure, enabling single-input operation. The dimensions of both the gripper and the re-entrant assembly were selected to ensure that the input and output regions remained within the microscope’s field of view. To facilitate direct observation of cell deformation, triangular pads were placed at the E, W, N, and S points to serve as attachment sites for individual cells (Fig. 2C). The triangular geometry allowed the arms to move into close proximity without premature contact. This design was microfabricated out of SU-8 using photolithography by a two-layer process described previously [11, 12].

### 2.2 Relations for estimating stretching force from mechanism deformation

In this section, we derive analytical relations to calculate the forces exerted on the mechanism by the cell during actuation. These forces are expressed as a function of the mechanism’s deformation. During stretching, the mechanism applies outward forces on the cell, and the cell, in turn, applies equal and opposite reaction forces on the mechanism. These reaction forces, denoted as *F*_1_ and *F*_2_, act in the East-West (E–W) and North-South (N–S) directions, respectively (Fig. 2D). To simplify the analysis, the geometric symmetry of the auxetic structure is used. A quadrant of the mechanism (Fig. 2E) is considered with symmetric boundary conditions. The forces *F*_1_ and *F*_2_ acting on the full structure are halved (Fig. 2F) when analyzing one quadrant. The procedure for calculating these forces is as follows:

- Obtain the expressions for the statically indeterminate boundary reactions using Castigliano’s second theorem.
- Obtain the expressions for the unknown displacements using Castigliano’s second theorem.
- Invert the expressions to obtain the forces in terms of the displacements. The variables used in the analysis are listed below:

*Q*_1_ : Reaction moment on the beam connecting points E and NE (North-East)

*Q*_2_ : Reaction moment on the beam connecting points N and NE

*F*_1_ : Force applied by the cell in the horizontal direction

*F*_2_ : Force applied by the cell in the vertical direction

*C* : Compliance matrix

*F*_out_ : Force applied on the auxetic by the gripper

*δ*_*x*_ : Horizontal displacement (between E and W)

*δ*_*y*_ : Vertical displacement (between N and S)

*R*_*E*_ : Vertical reaction at E

*R*_*N*_ : Horizontal reaction at N

*M*_*E*_ : Reaction moment at E (ccw)

*M*_*N*_ : Reaction moment at N (cw)

*F*_*A*_ : Axial force at a section

*F*_*T*_ : Transverse force at a section

By using the force balance in the horizontal and vertical directions and moment balance about E, we obtain relations between the reaction forces and moments in (Fig. 2F):

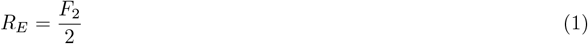

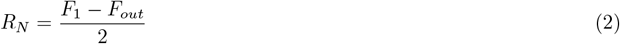

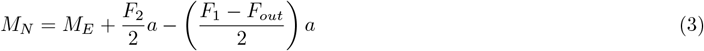

These expressions cannot solve all the reaction forces and moments because the system is statically indeterminate. To solve all the reaction forces and moments, we apply Castigliano’s second theorem, which requires the total strain energy as a function of all the external forces acting on the system.

The reaction moment at any arbitrary section can be obtained by taking moment balance about that section (Fig. 2G):

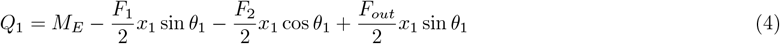

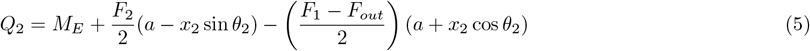

Using the Euler-Bernoulli Beam Theory, the strain energy expressions are:

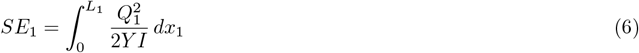

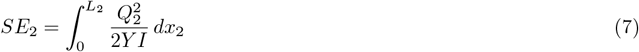

The total strain energy of the auxetic structure is:

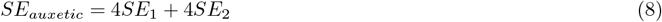

We employ the spring-lever model [14] to compute the strain energy of our gripper mechanism (Fig. 2H). Since the gripper is symmetric, we have divided it into two halves (Fig. 2I) and modeled each half using the spring-lever model (Fig. 2J and K). To determine the total strain energy of the entire gripper, we calculate the energy for one half and then double it.

The variables involved in the spring-lever model are as follows:

*k*_*ci*_ : Input port spring stiffness

*k*_*co*_ : Output port spring stiffness

*n* : Geometric amplification factor

*F*_*out*_ : Output force exerted by the gripper

*F*_*k*_ : Reaction force at the fixed support

*F*_*in*_ : Input force given by the actuator

*δ*_*in*_ : Displacement at the input point of the mechanism

The total strain energy of one half of the gripper is:

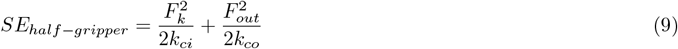

Since the system is in equilibrium, balancing the moment about an axis passing through the lever gives:

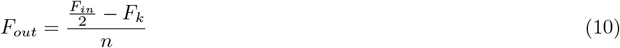

The total strain energy of the mechanism can now be written as the sum of the auxetic and gripper strain energies:

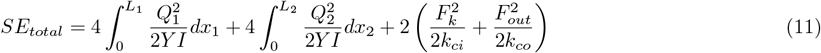

Equation (11) contains two unknowns: the moment *M*_*E*_, which appears in both *Q*_1_ and *Q*_2_, and the reaction force *F*_*k*_. Due to symmetry, the rotation at point E is zero. Additionally, since *F*_*k*_ is a reaction force, the displacement at the point where it acts is zero. These boundary conditions are applied using Castigliano’s second theorem:

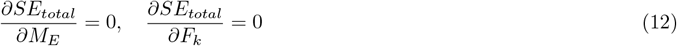

Castigliano’s second theorem is applied again to obtain expressions for the displacements in terms of the applied forces:

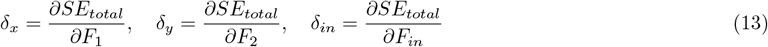

This leads to a linear relationship:

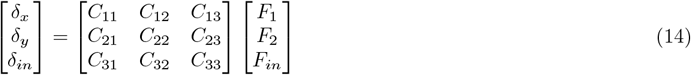

Finally, taking the inverse of the compliance matrix *C*:

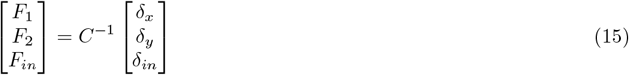

Using Eq. (15), the force exerted on the cell and the actuation force on the mechanism can be computed based on the horizontal (E–W), vertical (N–S), and input point displacements. This expression enables the determination of both the cell force and the input force from the displacements in the E–W, N–S, and input directions.

### 2.3 Cell attachment on the micromechanism and biaxial stretching

U87-MG human glioblastoma cells were selected for biaxial stretching experiments due to their strong adherence and frequent use in mechanobiological studies. The microfabricated mechanisms were sterilized using ultraviolet exposure inside a biosafety cabinet. To enhance cellular attachment, the mechanisms were coated with 50 *µ*g/mL of Collagen I (rat tail, Gibco, Cat. No. A10483-01). Cells were cultured in Dulbecco’s Modified Eagle Medium (DMEM; HiMedia, Cat. No. AT007), supplemented with 10% fetal bovine serum (FBS; Gibco, Cat. No. A3160801, Origin: Brazil), under standard conditions. After detachment via trypsinization, the cells were seeded directly onto the collagen-coated devices placed in 35 mm glass-bottom culture dishes (Gene X Biosciences, Cat. No. 8128-200) and incubated overnight to allow proper adhesion.

To actuate the micromechanism, the coverslip containing the device was mounted on a confocal microscope (Olympus IX83) [12]. A glass micropipette, fabricated using a pipette puller (Narishige PC-100) and mounted on an XYZ micromanipulator (Narishige, Co PC-100), served as a probe for actuation. The entire mechanism was submerged in media during cell stretching. Upon actuation, the displacements between the east-west ends (Δ*x*), north-south ends (Δ*y*), and the input point (Δ_in_) were quantified from bright-field microscopy images using ImageJ. Detailed experimental procedure is available in our previous publication [12].

## 3 Results and Discussion

### 3.1 Validating force relations by stretching extension springs

The force-displacement relationship, Eq. (15), was first verified on a macro-scale prototype of the design. The prototype was fabricated from 6 mm-thick acrylic sheets using laser cutting with an overall size of 650*×*650 mm and a minimum in-plane beam width of 4 mm. This device was used to stretch extension springs connected between the E-W and N-S points (Fig. 3) and the stiffness of the springs were estimated using the displacements and Eq. (15). These estimates were compared with spring stiffness estimated by hanging known weights.

**Figure 3:**
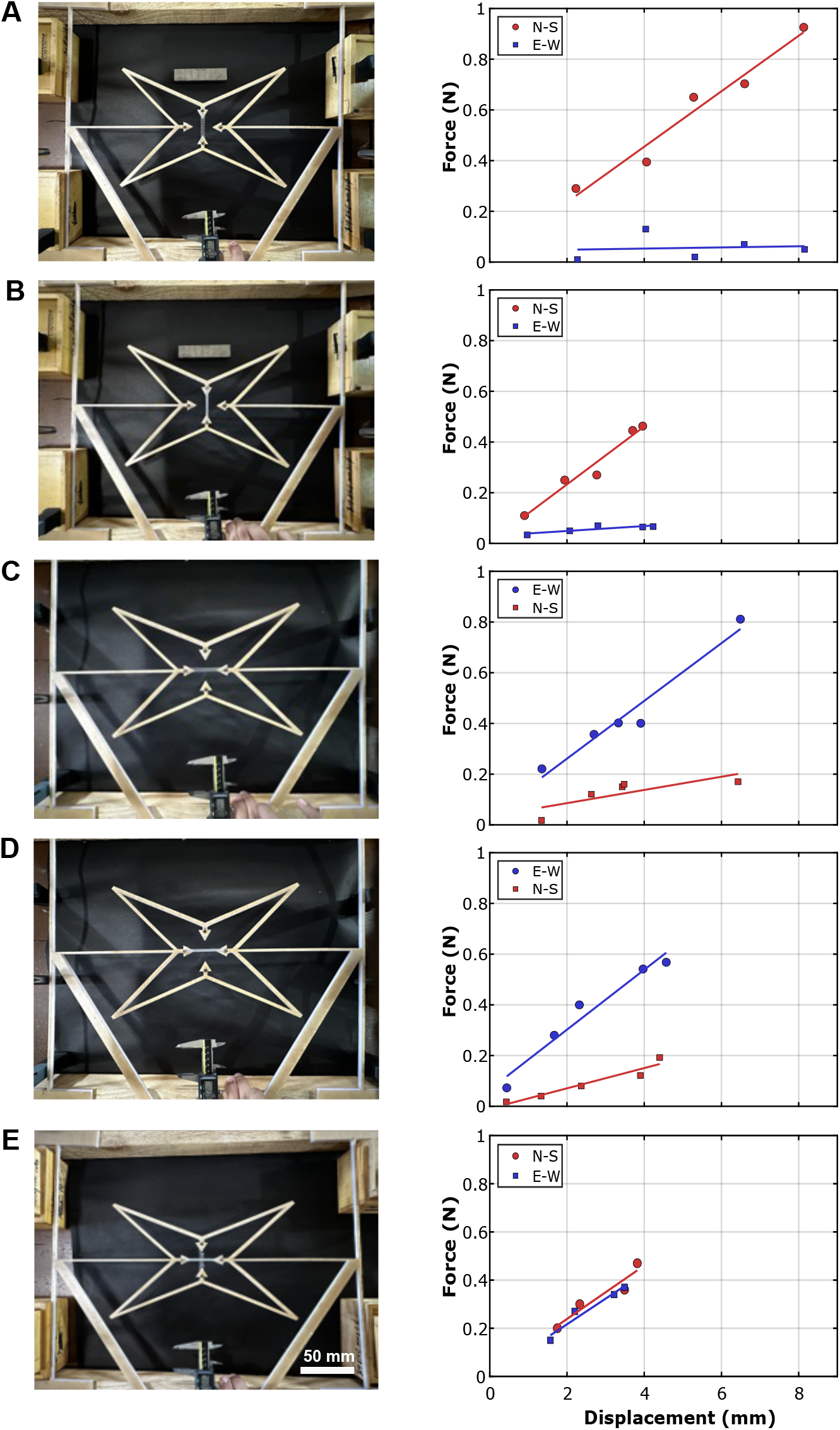
Estimating spring stiffness using the macro-scale prototype. Left column shows the experimental setup and right column shows the corresponding force-displacement data between E-W and N-S. (A) Spring A between N-S, (B) Spring B between N-S, (C) Spring A between E-W, (D) Spring B between E-W and (E) Spring A between E-W and Spring B between N-S

Two extension springs, designated A and B, were used in the experiments. By hanging known weights their stiffness were estimated as 108 N/m and 117 N/m respectively. By using these springs, a total of five stretching experiments were performed. In the first four, either Spring A or Spring B was connected to the mechanism in the E–W or N–S direction. In the fifth experiment, both springs were connected simultaneously—Spring A along the N–S direction and Spring B along the E–W direction.

The mechanism was actuated at the input point and the displacements at the input, E, W, N and S points were measured using image processing (ImageJ by National Institute of Health). Forces were obtained using Eq. (15). The parameters for the spring-lever model of the gripper, *k*_*ci*_, *k*_*co*_, and *n*, were obtained from finite element analysis (FEA) using ANSYS. The elastic modulus of acrylic required for FEA was estimated to be 3.15 GPa from three-point bending tests. The stiffness between E–W and N–S points was calculated by fitting a straight line to the force–displacement data (*F*_1_ vs. *δ*_*x*_ and *F*_2_ vs. *δ*_*y*_). These estimated stiffness values were then compared with the known spring constants. Table 1 summarizes the actual and estimated stiffness values for the five spring configurations. We note that the highest percentage error is around 9% whereas the average error is around 4%. The stiffness in the direction orthogonal to the spring, in case of the first four configurations, is typically one or two orders of magnitude lower than the spring stiffness further substantiating the accuracy of our force-displacement relationship, Eq. (15).

**Table 1:**
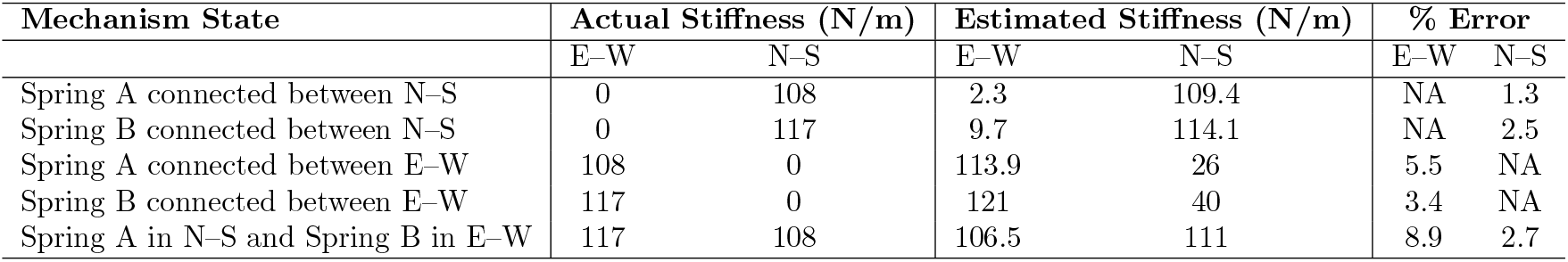
Comparison between actual and estimated stiffness values for different spring configurations. Stiffness values in the east–west (E–W) and north–south (N–S) directions are presented alongside the percentage error for each case. The estimated values are obtained using the analytical formulation given by Eq. (15).

| Mechanism State | Actual Stiffness (N/m) |  | Estimated Stiffness (N/m) |  | % Error |  |
| --- | --- | --- | --- | --- | --- | --- |
|  | E–W | N–S | E–W | N–S | E–W | N–S |
| Spring A connected between N–S | 0 | 108 | 2.3 | 109.4 | NA | 1.3 |
| Spring B connected between N–S | 0 | 117 | 9.7 | 114.1 | NA | 2.5 |
| Spring A connected between E–W | 108 | 0 | 113.9 | 26 | 5.5 | NA |
| Spring B connected between E–W | 117 | 0 | 121 | 40 | 3.4 | NA |
| Spring A in N–S and Spring B in E–W | 117 | 108 | 106.5 | 111 | 8.9 | 2.7 |

### 3.2 Validating force relations by stretching PDMS membranes

To further validate the force-displacement relations, Eq. (15), we characterized PDMS membranes using the macroscale prototype. The membranes were fabricated by mixing the polymer and crosslinker at a 20:1 weight ratio, followed by degassing to remove entrapped air. The mixture was cast into molds and cured at 40°C for 8 h. Square specimens with dimensions of 50 mm *×* 50 mm and a thickness of 1 mm were prepared, and surface markers were applied to enable strain measurement.

The specimens were mounted onto the mechanism using paper clips, and biaxial deformation was applied via a linear actuator positioned at the input point (Fig. 4A). Displacements in the E-W, N-S, and input point were measured using image analysis in ImageJ. These displacement values were substituted into Eq. (15) to calculate the forces acting on the membrane, following the same methodology used in the spring-based validation. The resulting force values were converted to first Piola–Kirchhoff stresses, *P*_*xx*_ and *P*_*yy*_, by dividing with the undeformed cross-sectional area. Stretches, *λ*_*xx*_ and *λ*_*yy*_, were computed based on the displacements of the surface markers (highlighted with magenta and green boxes in Fig. 4A). A neo-Hookean material model was fitted to the stress–stretch data (Fig. 4C and D) and the shear modulus was obtained as 126.7 *±* 6.6 kPa for n=3 samples. A previous study using standard biaxial testing machines had reported a shear modulus of 120 kPa [15]. Hence, the error in estimation of shear modulus of PDMS membranes using our mechanism was 5.6% .

**Figure 4:**
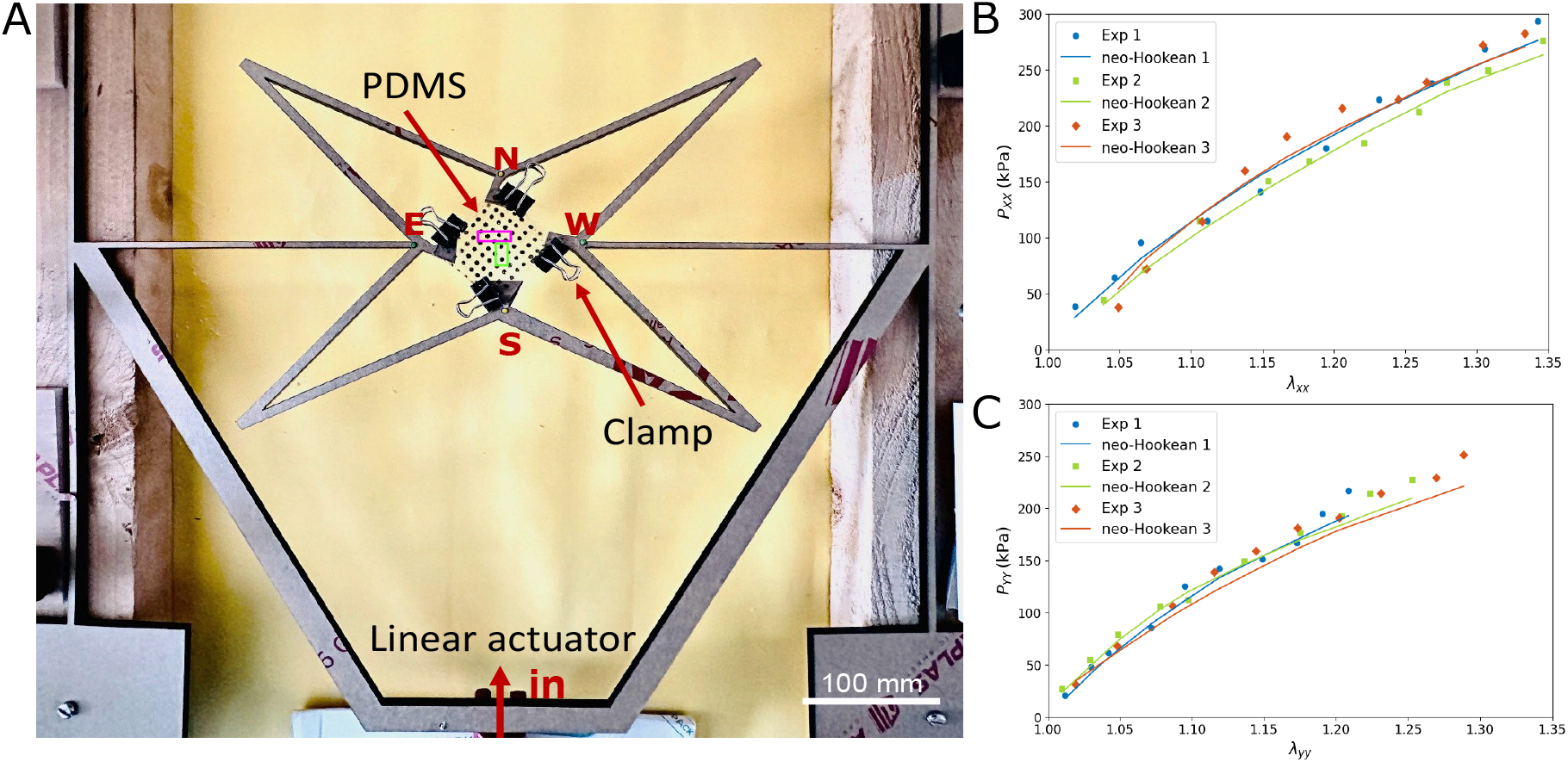
Biaxial stretching of PDMS membranes (A) Experimental setup. PDMS membrane is clamped on the stretcher using paper clips and the stretcher is actuated by a linear actuator at the input point. Markers used for measuring stretch in the x-direction (magenta box) and y-direction (green box) are highlighted. (B,C) First Piola-Kirchoff stress vs stretch in the x and y directions respectively. Circular markers are the measurements from three samples and the curves are the Neo-Hookean material model fitted to the experimental data.

While the motivation and focus of our work was to develop a micromechanism for biaxial stretching of single cells, we note that the macroscale prototype could be used for biaxial testing of tissues, films, polymer membranes and soft materials. In comparison to instruments such as the Biaxial Material Testing System (BiMaTS) [16], multi-functional biaxial stretching (BAXS) platform [17] and other commercial biaxial testers, which employ precision actuators with high cost, the present device offers a low-cost alternative that can be deployed without substantial infrastructure. The system’s portability also enables on-site mechanical testing immediately following tissue excision, eliminating transport-related delays and mechanical degradation [18]. This feature supports applications in tissue mechanics where immediate characterization is necessary, particularly outside dedicated laboratory environments.

### 3.3 Validating force relations on the micromechanism

Next, we validated the force-displacement relations, Eq. (15), on a microscale version of our design. The micromechanism was actuated using an optical-fiber probe capable of measuring the force that it exerts [19, 20]. Sequential images captured during actuation show the undeformed (Fig. 5A), intermediate (Fig. 5B), and the final (Fig. 5C) deformed configurations. It should be noted that cells are not attached at the E, W, N, and S points during this experiment, resulting in zero external forces at these locations. The force vs. displacement at the input point measured using the optical-fiber force transducer probe is shown in Fig. 5D. From this force-displacement data, the stiffness at the input point was estimated as 0.52 N/m. The displacements at the E-W, N-S and input points during this actuation was measured from the captured images using ImageJ. To calculate the forces from these displacements using Eq. (15), the elastic modulus of SU-8 (material of the micromechanism) is required. Since the *F*_*in*_ can be estimated from *δ*_*in*_ using the stiffness of the input point measured using optical probe (Fig. 5D), we used that to calibrate the elastic modulus of SU-8. We found that at 2.1 GPa, the error in estimation of *F*_*in*_ was *<* 5% (Table 2). This is well within the 0.9 -7.4 GPa range reported previously [21]. For all the configurations of the mechanism, *F*_*x*_ and *F*_*y*_ is close to zero, *<* 2% of *F*_*in*_ (Table 2), further demonstrating the accuracy of the force-displacement relations given in Eq. (15).

**Table 2:** Comparison of analytical and experimental input forces for the microfabricated mechanism. Displacements in the east–west (Δ*x*), north–south (Δ*y*), and input (Δ_in_) directions are used as inputs to the analytical model defined in Eq. (15). The resulting force components along the east–west (*F*_*x*_) and north–south (*F*_*y*_) directions, as well as the input force (*F*_in_), are calculated analytically. The analytical value of *F*_in_ is compared with the force measured experimentally at the input point.

| $\Delta x$ ($\mu\text{m}$) | $\Delta y$ ($\mu\text{m}$) | $\Delta_{in}$ ($\mu\text{m}$) | $F_x$ ($\mu\text{N}$) | $F_y$ ($\mu\text{N}$) | $F_{in}$ (Analytical) ($\mu\text{N}$) | $F_{in}$ (Experimental) ($\mu\text{N}$) | % Error |
| --- | --- | --- | --- | --- | --- | --- | --- |
| 2.98 | 2.95 | 5.00 | 0.03 | 0.01 | 2.58 | 2.57 | 0.38 |
| 4.30 | 4.15 | 7.50 | 0.08 | 0.05 | 3.91 | 4.05 | 3.45 |
| 5.95 | 5.91 | 10.00 | 0.07 | 0.01 | 5.20 | 5.45 | 4.58 |
| 7.51 | 7.46 | 12.50 | 0.05 | 0.02 | 6.43 | 6.72 | 4.31 |

**Figure 5:**
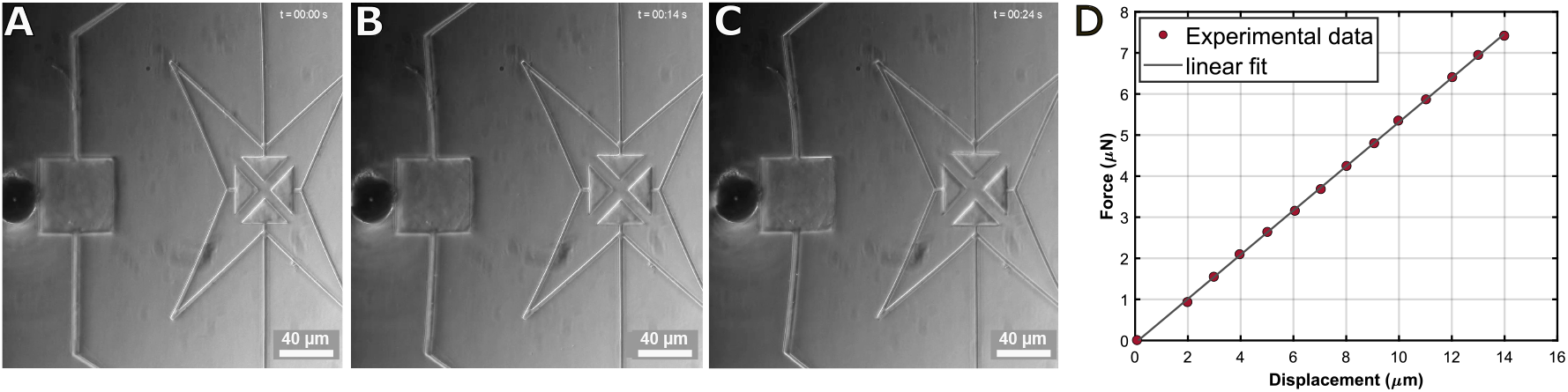
Actuating the micromechanism using an optical-fiber probe (A) Undeformed configuration. (B) Low biaxial stretch. (C) High biaxial stretch. (D) Force–displacement response at the input point of the mechanism measured using the optical-fiber probe

### 3.4 Force resolution

Since the forces are estimated from the displacements, Eq. (15), the errors in forces are due to the errors in displacements. The stiffness matrix, *K* = *C*^*−*1^, functions like a scaling factor that either amplifies or reduces displacement errors. Since we are only interested in the cell stretching forces (*F*_1_ and *F*_2_), and not the input force (*F*_*in*_), we can further reduce Eq. (15) to

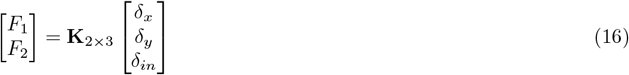

where

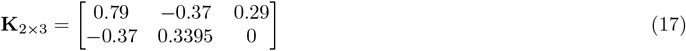

for our micromechanism (unit -N/m). It may be noted that *F*_2_ is decoupled from *δ*_*in*_ (***K***_2*×*3_(2, 3) = 0) because the gripper is connected at E and W and not at N and S. The displacement resolution *d* is given by the diffraction-limited resolution of the imaging system:

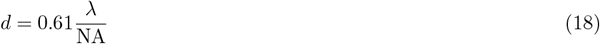

where *λ* =400 nm is the laser wavelength and NA = 0.75 is the numerical aperture of the objective lens (20X). By substituting these values we obtained the displacement resolution as 0.325 *µ*m. For obtaining the force resolution we performed Monte-Carlo simulations. *δ*_*x*_, *δ*_*y*_ and *δ*_*in*_ were randomly drawn from uniform distributions between [-d/2 d/2] and the corresponding forces, *F*_1_ and *F*_2_, were calculated using Eq. (16). A total of 100000 simulations were performed. Since the distributions of *F*_1_ and *F*_2_ are a weighed sum of three and two uniform distributions respectively, Eq. (17), the resulting distributions are bell-shaped for *F*_1_ and triangular for *F*_2_. From the cumulative distribution of *F*_1_ and *F*_2_ we calculated the range, centered around the mean, that contained 90% of the data. The full width of this range was deemed as the force resolution, which was 0.29 *µ*N and 0.16 *µ*N for *F*_1_ and *F*_2_ respectively.

To the best of our knowledge, there is only one previous report of a mechanisms-based single-cell biaxial stretcher [22]. The MEMS device reported in that study consisted of a comb-drive actuated platform with four quadrants moving in orthogonal directions. For cells to remain viable during stretching, they must be submerged in a solution containing ions, which hinders the comb drive actuator, thereby compromising the performance of the device. Hence, biaxial stretching of cells was not demonstrated using this device.

In the absence of a biaxial manipulation for direct comparison, we examine micromechanisms and MEMS devices developed for simpler uniaxial manipulation modes such as squeezing [23, 24], poking [25] and stretching [26, 27]. Several of them report resolutions in tens of nNs primarily through a large footprint, size of 2-10 millimeters in comparison to our device with a size of 500 *µ*m. The longer beam lengths reduce the device stiffness thereby improving resolution, which however limits scalability and constrains integration into lab-on-a-chip systems. All devices, except [23] which uses SU-8, are made from silicon, which increases costs and restricts imaging on inverted microscopes—–commonly used in biological studies—–due to silicon’s opacity. Among, other traditional single-cell manipulation techniques (not based on micromechanisms), OT can achieve pN force resolution [28]. However, it is an expensive and tedious technique with limited manipulation modes, mostly uniaxial stretching. Further, the low magnitude of forces achieved in OT (few nNs [3]), restrict the magnitude of cell deformation possible with this technique. In case of AFM, the enhanced force resolution (sub-nN) is due to the superior displacement resolution (sub-nm) achieved using the Position-Sensitive Photodiode (PSPD). However, PSPD limits AFM cell manipulations to only indentations. Hence, the better force resolution of these techniques are achieved at the cost of experimental complexity and limited manipulation modes. While this tradeoff maybe suitable for single-molecule studies (forces in the range of pNs), in case of whole-cell deformation studies (forces in the range 0.1 - 10 *µ*N), a micromechanisms-based strategy with a lower force resolution but higher manipulation versatility and experimental simplicity might be advantageous.

Force resolution in our mechanisms is constrained by the displacement resolution which is determined by the optics of the imaging system and therefore the force resolution can be improved by enhancing the displacement resolution. This could be achieved through high-resolution objectives (NA = 1.4) and displacement tracking techniques at sub-pixel resolution in bright field [24] or fluorescent images [29].

### 3.5 Biaxial stretching of a single cell and cell force measurement

Finally, we present the estimation of anisotropy of single cells. Biaxial stretching of a single cell using our mechanism is shown in Fig. 6A-C. The cell is firmly attached to all four triangular pads, as confirmed by live imaging under confocal microscopy. The stretching force was estimated from the mechanism deformation, which was converted to First Piola-Kirchoff stresses by dividing by the cross-sectional area. For this, the thickness of the cell was assumed to be 5 *µ*m. The stretches in the horizontal (*λ*_*xx*_) and vertical (*λ*_*yy*_) directions of the cell were measured from the images. A 2D Fung model with the following strain energy density function was fit to the stress-stretch data (Fig. 6D and E).

**Figure 6:**
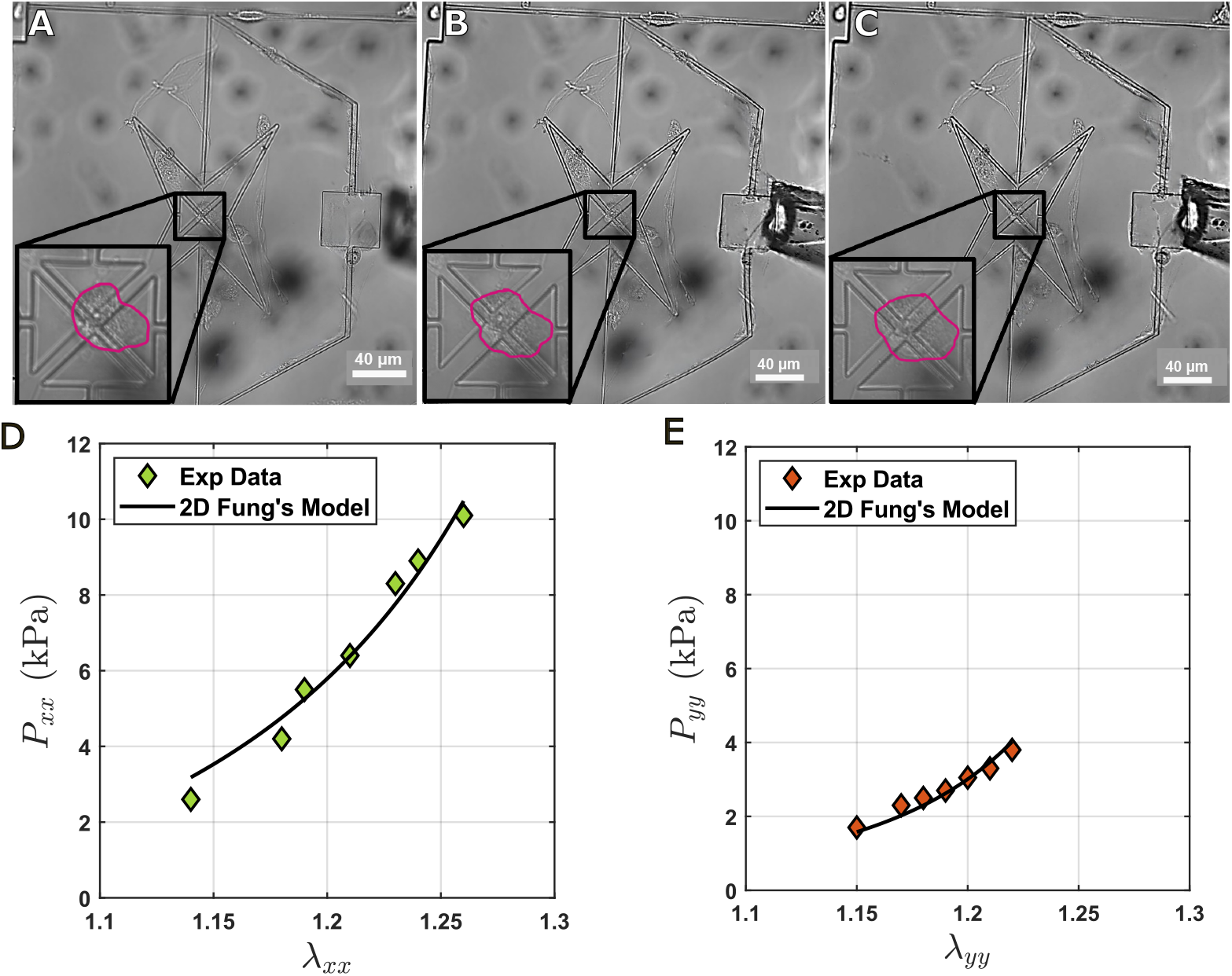
Estimating cellular anisotropy. (A-C) Brightfield images of biaxial stretching. (A) unactuated (B) Low biaxial stretching and (C) High biaxial stretching. First Piola-Kirchoff stress vs stretch in (D) E-W (x) (E) N-S (y) directions. Markers are the experimental measurements and curves are the 2D Fung model fit to the data

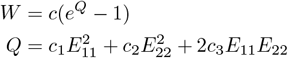

The material parameters obtained from fitting this model to biaxial cell stretching data (Table 3) indicate cellular anisotropy.

**Table 3:**
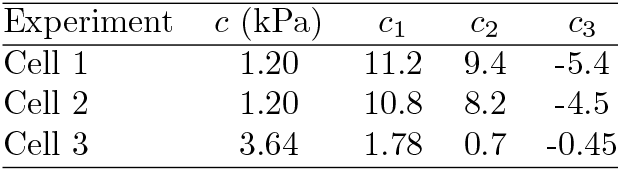
Material parameters obtained by fitting the 2D Fung model to biaxial cell stretching data. The parameters *c*_1_ and *c*_2_ reflect stiffness in the E-W and N-S directions, respectively, while *c*_3_ indicates coupling between these directions. The parameter *c* is a scaling constant of the strain energy function.

| Experiment | $c$ (kPa) | $c_1$ | $c_2$ | $c_3$ |
| --- | --- | --- | --- | --- |
| Cell 1 | 1.20 | 11.2 | 9.4 | -5.4 |
| Cell 2 | 1.20 | 10.8 | 8.2 | -4.5 |
| Cell 3 | 3.64 | 1.78 | 0.7 | -0.45 |

The values of *c*_1_ and *c*_2_, which represent the stiffness contributions in the *x* and *y* directions, respectively, are distinct (e.g., *c*_1_ = 11.2 vs. *c*_2_ = 9.4 for the first cell), indicating anisotropy. Previous studies have shown that cells align their cytoskeleton in the direction of higher substrate stiffness resulting in higher cellular stiffness in that direction [30]. For our mechanism, the stiffness is higher in the *x* -direction (E–W) (0.3 N/m) than in the *y* -direction (N–S) (0.15 N/m) because the gripper is connected to the E and W ends. Hence, our observation of higher cellular stiffness between E-W in comparison to N-S, *c*_1_ *> c*_2_ for all cells, is consistent with previous results. Most previous studies of biaxial stretching have cultured cells on flexible membranes and stretched the substrate biaxially [31, 32, 33]. These are mechanobiology studies, wherein the focus is to understand the biological effects of biaxial stretching on cells and cell populations and not mechanical characterization. To the best of our knowledge, the only studies that has measured cellular anisotropy using biaxial stretching are [34, 35]. They developed a biaxial substrate stretcher with polyacrylamide gels containing fluorescent beads and characterized the anisotropy of Vascular Smooth Muscle Cells by fitting a Holzapfel–Gasser–Ogden type strain energy density function. While the biaxial stretching method, cell type (Vascular Smooth Muscle Cells) and material model used in these studies are different from ours, they have also observed cellular anisotropy similar to our study.

In the absence of other biaxial stretching studies on U87-MG cells, we benchmarked our results with studies that have measured the Young’s modulus of these cells. For obtaining the Young’s modulus from the 2D Fung model we used the following expression for a uniaxial stretching experiment (*σ*_22_ = 0)

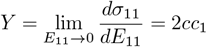

From the material parameters fitted to our experiments (Tab. 3) we obtained Y = 21.92 *±* 7.78 kPa. A previous study using AFM on U87-MG cells had reported a Young’s modulus of 24.62 *±* 2.05 kPa [36] (10.8% error). Hence, our micromechanism is able to recapitulate the cellular anisotropy and Young’s modulus from previous studies.

## 4 Conclusion

We have demonstrated a micro biaxial stretcher for estimating the anisotropy of single cells. Since our design is based on a compliant mechanism wherein the motion is transmitted via deformation of the mechanism, the forces acting on the cells can be computed from the mechanism deformation. Hence, unlike MEMS techniques, other force sensing methods such as capacitance or piezo-resistance need not be integrated into the device, which simplifies the fabrication and experimentation. An analytical relationship between the cell stretching forces and mechanism deformations was derived using Euler-Bernoulli beam theory and Castigliano’s theorems. These equations were verified both at macro- and micro-scales. While at the macro-scale we used extension springs and PDMS membranes, at the microscale we used an optical fiber-based force transducer. By using Monte-Carlo simulations were estimated the force resolution of the device as 0.29 *µ*N and 0.16 *µ*N in the horizontal vertical directions respectively. Finally, we estimated the anisotropy of glioblastoma cells by fitting a 2D Fung model to the biaxial stress-stretch data. Our results show a higher stiffness in the horizontal direction in comparison to the vertical direction, which could be because the mechanism is stiffer along the horizontal direction. Our device enables direct force measurement and mechanical characterization of single cells without integrated sensing elements. It combines compliant design, image-based tracking, and analytical modeling in a compact and scalable format.

In the future, we aim to expand this platform to include single-cell twisting and shearing by developing additional compliant mechanisms (Fig. 1C and D). By integrating uniaxial, biaxial, shear, and twist modules onto a single chip, we can create a multi-modal lab-on-a-chip platform for single-cell manipulation. Since this platform supports high-resolution imaging, the alterations in the shapes of the cell and its organelles during manipulation could be tracked enabling further physical characterization using morphology-based techniques [37, 38, 39]. In summary, the mechanisms-based framework enables systematic studies of how individual cells respond to different mechanical inputs and provide a foundation for high-throughput mechanobiological screening, disease modeling, and therapeutic testing.

## Acknowledgments

Authors thank Indian Nanoelectronics Users Program for device fabrication and Dr. Anju R Babu, National Institute of Technology Rourkela, India for sharing biaxial testing data of PDMS membranes; H.M. is grateful for Ministry of Education funding during PhD and S.B. thanks Science and Engineering Research Board, Government of India (grant number - SRG/2020/001555) for funding.

## References

[1] I. Sokolov, M. E. Dokukin, N. V. Guz, Methods 2013, 60, 2 202.

[2] R. M. Hochmuth, Journal of biomechanics 2000, 33, 1 15.

[3] H. Zhang, K.-K. Liu, Journal of the Royal Society interface 2008, 5, 24 671.

[4] Review of Scientific Instruments 2006, 77, 5.

[5] Y. Zhang, F. Wei, Y.-C. Poh, Q. Jia, J. Chen, J. Chen, J. Luo, W. Yao, W. Zhou, W. Huang, et al., Nature protocols 2017, 12, 7 1437.

[6] S. Schürmann, S. Wagner, S. Herlitze, C. Fischer, S. Gumbrecht, A. Wirth-Hücking, G. Prölß, L. Lautscham, B. Fabry, W. Goldmann, et al., Biosensors and bioelectronics 2016, 81 363.

[7] P. H. Wu, D. R. B. Aroush, A. Asnacios, W. C. Chen, M. E. Dokukin, B. L. Doss, P. Durand-Smet, A. Ekpenyong, J. Guck, N. V. Guz, P. A. Janmey, J. S. Lee, N. M. Moore, A. Ott, Y. C. Poh, R. Ros, M. Sander, I. Sokolov, J. R. Staunton, N. Wang, G. Whyte, D. Wirtz, Nature Methods 2018, 1–8.

[8] S. Kollimada, S. Balakrishnan, C. K. Malhi, S. R. Raju, M. Suma, S. Das, G. Ananthasuresh, Journal of Micro-Bio Robotics 2017, 13 27.

[9] S. A. Kollimada, S. Khan, S. Balakrishnan, S. Raju, S. Matad, G. K. Ananthasuresh, International Conference on Manipulation, Automation and Robotics at Small Scales, MARSS 2017 - Proceedings 2017, 1–6.

[10] N. S. Fartyal, H. Marwah, S. Balakrishnan, In International Design Engineering Technical Conferences and Computers and Information in Engineering Conference, volume 85444. American Society of Mechanical Engineers, 2021 V08AT08A009.

[11] H. Marwah, R. Nautiyal, N. S. Fartyal, S. Balakrishnan, In 2023 International Conference on Manipulation, Automation and Robotics at Small Scales (MARSS). IEEE, ISBN 979-8-3503-3039-7, 2023 1–6, URL https://ieeexplore.ieee.org/document/10294156/.

[12] H. Marwah, N. Fartyal, H. Bhatt, R. Nautiyal, S. Balakrishnan, Journal of Micro and Bio Robotics 2024, 20, 2 14.

[13] K. E. Evans, Endeavour 1991, 15, 4 170.

[14] S. Hegde, G. Ananthasuresh, ASME 2010.

[15] A. Babu, N. Gundiah, Experimental Mechanics 2014, 54 1177.

[16] A. Corti, T. Shameen, S. Sharma, A. De Paolis, L. Cardoso, HardwareX 2022, 12 e00333.

[17] D. Tremblay, C. M. Cuerrier, L. Andrzejewski, E. R. O’Brien, A. E. Pelling, Journal of visualized experiments: JoVE 2014,, 88 51454.

[18] O. Friedrich, A.-L. Merten, D. Schneidereit, Y. Guo, S. Schürmann, B. Martinac, Frontiers in Bioengineering and Biotechnology 2019, 7 55.

[19] C. Kalelkar, P. A. Pullarkat, et al., Review of Scientific Instruments 2013, 84, 10.

[20] S. Dubey, S. Veer, R. S. Rao, C. Kalelkar, P. A. Pullarkat, Journal of Physics: Condensed Matter 2020, 33, 8 084003.

[21] T. Xu, J. H. Yoo, S. Babu, S. Roy, J. B. Lee, H. Lu, Journal of Micromechanics and Microengineering 2016, 26, 10 0.

[22] N. Scuor, P. Gallina, H. V. Panchawagh, R. L. Mahajan, O. Sbaizero, V. Sergo, Biomedical Microdevices 2006, 8,3 239.

[23] S. D. Bhargav, N. Jorapur, G. Ananthasuresh, Mechanism and Machine Theory 2015, 91 258.

[24] B. Barazani, S. Warnat, A. Fine, T. Hubbard, Journal of Micromechanics and Microengineering 2016, 27, 2 025002.

[25] S. Yang, M. T. A. Saif, Acta Biomaterialia 2007, 3, 1 77.

[26] D. B. Serrell, T. L. Oreskovic, A. J. Slifka, R. L. Mahajan, D. S. Finch, Biomedical microdevices 2007, 9 267.

[27] V. Mukundan, B. L. Pruitt, Journal of Microelectromechanical Systems 2009, 18, 2 405.

[28] C. J. Bustamante, Y. R. Chemla, S. Liu, M. D. Wang, Nature Reviews Methods Primers 2021, 1, 1.

[29] M. K. Cheezum, W. F. Walker, W. H. Guilford, Biophysical Journal 2001, 81, 4 2378.

[30] A. Saez, M. Ghibaudo, A. Buguin, P. Silberzan, B. Ladoux, Proceedings of the National Academy of Sciences 2007, 104, 20 8281.

[31] C. Simmons, J. Sim, P. Baechtold, A. Gonzalez, C. Chung, N. Borghi, B. Pruitt, Journal of Micromechanics and Microengineering 2011, 21, 5 054016.

[32] D. Tremblay, S. Chagnon-Lessard, M. Mirzaei, A. E. Pelling, M. Godin, Biotechnology letters 2014, 36 657.

[33] J. Imsirovic, T. J. Wellman, J. R. Mondoñedo, E. Bartolák-Suki, B. Suki, PLoS One 2015, 10, 10 e0140283.

[34] Z. Win, J. M. Buksa, K. E. Steucke, G. W. G. Luxton, V. H. Barocas, P. W. Alford, Journal of biomechanical engineering 2017, 139, 7 071006.

[35] Z. Win, J. M. Buksa, P. W. Alford, Biophysical Journal 2018, 115, 10 2044.

[36] N. Masud, M. H. H. Hasib, B. Ibironke, C. Block, J. Hughes, A. Ekpenyong, A. Sarkar, Scientific Reports 2025, 15, 1 1.

[37] S. Balakrishnan, S. S. Mathad, G. Sharma, S. R. Raju, U. B. Reddy, S. Das, G. Ananthasuresh, Biophysical Journal 2019, 116, 7 1328.

[38] S. Balakrishnan, S. Raju, A. Barua, R. P. Pradeep, G. K. Ananthasuresh, Biophysical Journal 2021, 120, 21 4698.

[39] A. Devulapally, V. Parekh, C. Pazhayidam George, S. Balakrishnan, Cells Tissues Organs 2024, 213, 2 96.

